# Can herbivore-induced volatiles protect plants by increasing the herbivores’ susceptibility to natural pathogens?

**DOI:** 10.1101/317560

**Authors:** Laila Gasmi, María Martínez-Solís, Ada Frattini, Meng Ye, María Carmen Collado, Ted C.J. Turlings, Matthias Erb, Salvador Herrero

## Abstract

In response to insect herbivory, plants mobilize various defenses. Defense responses include the release of herbivore-induced plant volatiles (HIPVs) that can serve as signals to alert undamaged tissues and to attract natural enemies of the herbivores. It has also been shown that some HIPVs can have a direct negative impact on herbivore survival, but it is not yet understood by what mechanism. Here we tested the hypothesis that exposure to HIPVs renders insects more susceptible to natural pathogens. Exposing caterpillars of the noctuid *Spodoptera exigua* to indole and linalool, but not exposure to (*Z*)-3-hexenyl acetate increased the susceptibility to its nucleopolyhedrovirus (SeMNPV). We also found that exposure to indole, but not exposure to linalool or (*Z*)-3-hexenyl acetate, increased the pathogenicity of *Bacillus thuringiensis*. Additional experiments revealed significant changes on gut microbiota composition after forty-eight hours of larval exposure to indole. Overall, these results provide evidences that certain HIPVs can strongly enhance the susceptibility of caterpillars to pathogens, possibly through effects on the insects’ gut microbiota. These findings suggest a novel mechanism by which HIPVs can protect plants from herbivorous insects.

## Introduction

Plants defend themselves against herbivores through the production of specific metabolites and proteins with toxic, repellent, or antinutritive properties (Jander & Howe, 2008). These defense compounds are either produced constitutively or induced in response to herbivore attack (Schoonhoven *et al*., 2005). Induction is mainly mediated by the insect feeding and leads to the activation of multiple signaling pathways that regulate the production of defensive proteins and metabolites (Mithöfer & Boland, 2012; War *et al*., 2012; Lazebnik *et al*., 2014). Herbivores exhibit multiple feeding styles (i.e. chewing, sucking) and differ in the levels of specialization to their host plants. Accordingly, the plant response can vary depending on the type of herbivore and can involve a combination of responses in case of multiple attacks. Plant defense responses can also be elicited by other herbivore-related factors such as oviposition by insects (Hilker & Fatouros, 2015; Veyrat *et al*., 2016) or even by the perception of volatiles emitted by neighboring plants in response to insect attack (Kim & Felton, 2013; Erb *et al*., 2015).

Plant-emitted volatiles represent a group of specialized metabolites that play an important role in the plant defense against herbivory. Attacked plants release herbivore-induced plant volatiles (HIPVs) which can act as priming signals (Heil & Silva Bueno, 2007; Frost *et al*., 2008; Erb *et al*., 2015) or attract herbivore natural enemies (Dicke & Sabelis, 1987; Turlings *et al*., 1990; De Moraes *et al*., 1998). HIPVs can also have direct benefits for the plant by repelling the herbivore or reducing its growth and survival in the plant (Maag *et al*., 2015). For instance, the Green leaf volatile (Z)-3-hexenol from infested neighbors plants was found to be converted to (Z)-3-hexenyl-vicianoside in tomato (*Solanum lycopersicum*), reducing survival and growth of *Spodoptera littura* caterpillars (Sugimoto *et al*., 2014). More recently, it has also been shown that the HIPV indole increases weight gain, but reduces food consumption and survival in *Spodoptera littoralis* (Veyrat *et al*., 2016).

In a multi-trophic context, the eventual outcome of the interaction between plant and herbivore is also modulated by pathogenic microbes, which is assumed to be due to direct as well as indirect effects of toxic phytochemicals on entomopathogen persistence and infectivity (Cory & Hoover, 2006; Shikano, 2017; Shikano *et al*., 2017). Although some authors have speculated about the possibility of plant promoting the action or abundance of microbial entomopathogens (Elliot *et al*., 2000) no much information is available about the impact of HIPVs on the pathogenicity of entomopathogens. So far, only few studies have reported the influence of certain plant volatiles on the conidial germination rates of entomopathogenic fungi (Brown *et al*., 1995; Lin *et al*., 2016). To test this, we investigated the effect of specific HIPVs on the pathogenicity of two types of entomopathogens that naturally infect the beet armyworm, *Spodoptera exigua*. Larval mortality due to the *Spodoptera exigua nucleopolyhedrovirus* (SeMNPV) and to *Bacillus thuringiensis* was measured during exposure of the insect to indole, linalool, or hexenyl-acetate. In addition, we evaluated the effect of these volatiles on insect cellular immunity and gut microbiota composition. The results reveal a novel indirect defensive role for HIPVs by enhancing the pathogenicity of entomopathogens.

## Results

### HIPVs effects on the susceptibility to viral and bacterial pathogens

Newly molted third instar *S. exigua* larvae were used to evaluate the effect of HIPVs on insect susceptibility to a sublethal dose of the SeMNPV. For that, the viral dose was set to produce about 10-20% of mortality under our experimental conditions. Larvae were reared on artificial diet in the presence of a 0,2 ml tube that either was empty (control), contained 4 mg of indole, or contained a 10% solution of linalool or hexenyl-acetate. The lid of each tube was punctured to allow the release of the volatile during the duration of the bioassay. Compared with the control conditions in the absence of HIPVs, a drastic increase in mortality (P<0.001) due to baculovirus infection was observed when larvae where reared in the presence of indole or linalool (Fig. 1A and S1). No effect on SeMNPV pathogenicity was observed when the larvae were exposed to hexenyl-acetate (Fig. 1A and S1). Significant synergistic interaction (P<0.001) was found between the SeMNPV virus and indole or linalool. At the SeMNPV dose that we used, no increase in virulence (measured as the mean of time to death by the viral infection) was observed for any of the HIPVs treatments (Fig. 1B and S1).

**Fig. 1.**
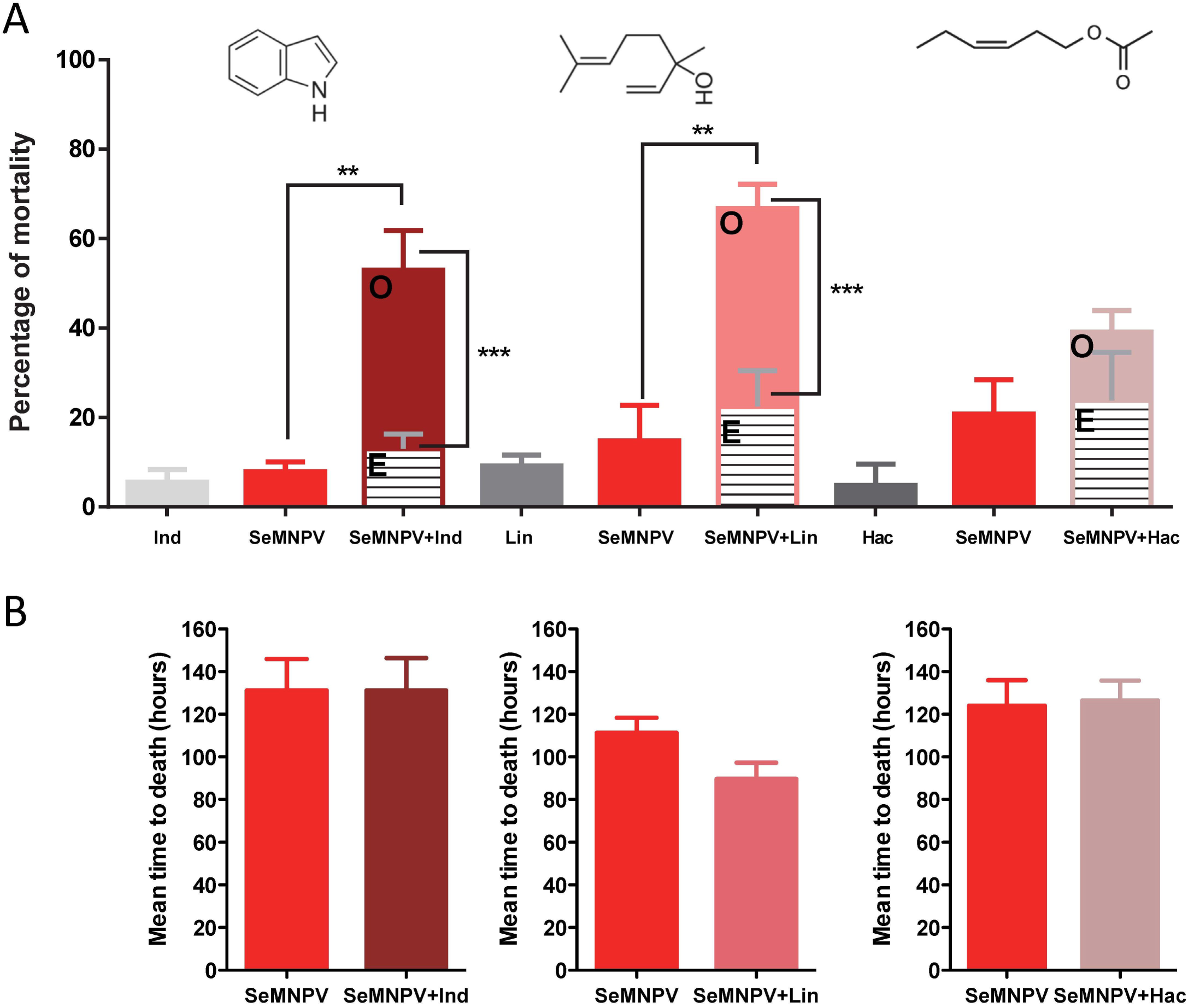
Effect of the tested HIPVs on the *S. exigua* susceptibility to SeMNPV infection (10^2^ OBs/larvae). Ind (indole 4 mg), Lin (linalool 10%), and Hac (hexenyl acetate 10%.) A) Percentage of larval mortality for the different combinations. Observed mortality (O) and Expected mortality (E) assuming the additive model. Statistic analyses were performed using the Newman-Keuls test to compare the mortalities and the Fisher’s exact test was used to check whether there is synergism or additive effect between the different treatments (GraphPad Prism5). ** refers to P-value<0.001 and *** refers P-value<0.0001 B) Mean time to death produced by baculovirus in the presence/absence of the corresponding HIPV. Values were statistical compared using student’s t-test.

The effect of exposure to the indole on the SeMNPV infectivity was also tested at a higher viral dose (producing about 80-90% mortality). Under these conditions, no additional increase in mortality was observed in the presence of indole, however a significant increase in virulence of the virus was found, with mortality occurring 20% earlier in the indole-exposed insects (Fig. S2A and S2B). In a more controlled environment where indole was released at a similar rate as produced by maize plants (50 ng/h) (Erb *et al*., 2015), we also observed a significant increase in baculovirus virulence (Fig. S2C and S2D).

We also evaluated the effects of HIPVs on the insect’s susceptibility to a bacterial pathogen (Fig. 2). Under our experimental conditions, mortality due to *B. thuringiensis* was not affected by exposure to linalool or (*Z*)-3-hexenyl acetate. However, a significant additive interaction (P<0.05) was found between *B. thuringiensis* and indole.

**Fig. 2.**
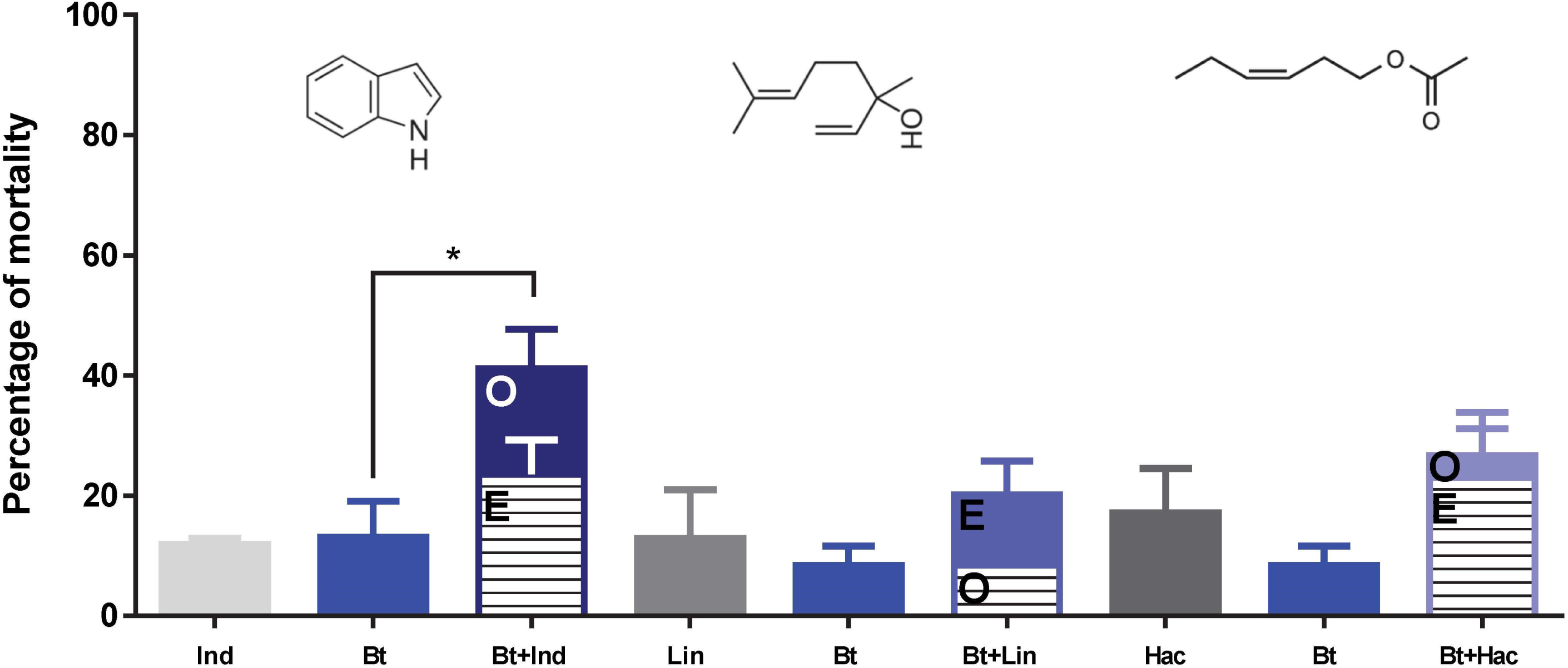
Effect of the tested HIPVs on the *S. exigua* susceptibility to *B. thuringiensis* (Xentari ^®^). Ind (indole 4 mg), Lin (linalool 0.1%) and Hac (hexenyl acetate 0.1%.). Percentage of larval mortality for the different combinations. Observed mortality (O) and Expected mortality (E) assuming the additive model. Statistic analyses were performed using the Newman-Keuls test to compare the mortalities and the Fisher’s exact test was used to check whether there is synergism or additive effect between the different treatments (GraphPad Prism5). * refers to P-value<0.05.

### Immune status of insects exposed to volatiles

To test if exposure to HIPVs affects the immunological status of *S. exigua*, we measured the levels of two enzymatic key markers of the cellular immunity in insects, phenoloxidase (PO) and phospholipase A2 (PLA2). PO is involved in the process of encapsulation and melanization (Cerenius & Soderhall, 2004), whereas the enzyme PLA2 activates the eicosanoid pathway involved in the cellular immunity in insects (Stanley *et al*., 2009). Several studies have shown that the inhibition of eicosanoids increases insect susceptibility to baculovirus (Stanley & Shapiro, 2009; Kim & Kim, 2011). PO activity was measured in the haemolymph of L3 larvae exposed to the different HIPVs for 24 and 48 hours. Compared to controls, the exposure had no effect on PO activity (Fig 3A). PLA2 activity was measured on the whole body extract of L3 larvae exposed to the three HIPVs for 24 and 48 hours (Fig 3B). Again, no effect on enzyme activity was observed for any of the treatments. These results suggest that the observed effect of HIPV exposure on the susceptibility to pathogens is unlikely to be mediated by changes in the cellular immunity.

**Fig. 3.**
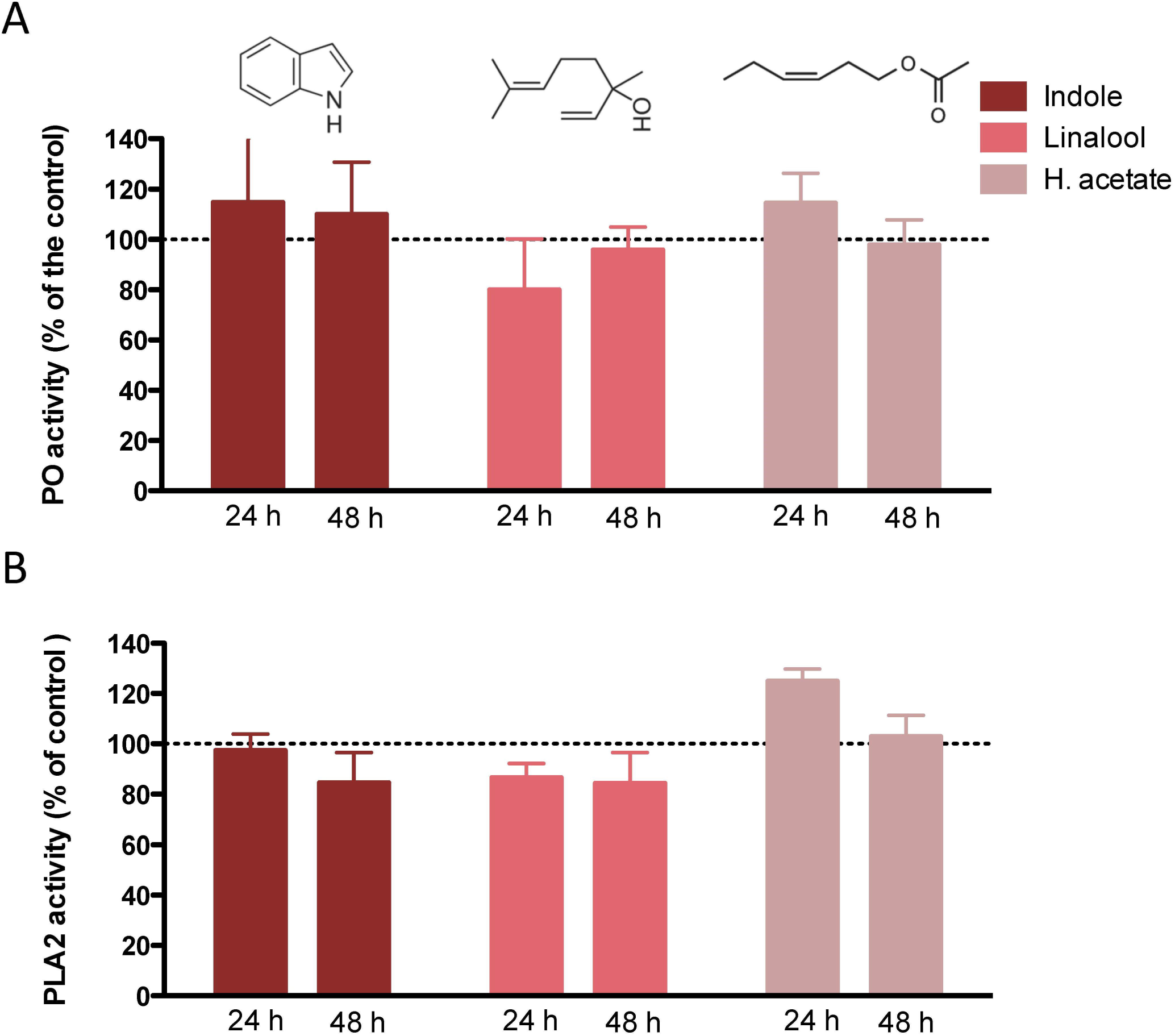
Effect of the tested HIPVs on two enzymatic markers of the cellular immunity of *S. exigua*. A) Relative Phenoloxidase activity in the haemolymph of insects exposed to selected volatiles at 24 and 48 hours after exposure. B) Relative PLA2 activity in the fat body of insects exposed to selected volatiles at 24 and 48 hours after exposure. For both markers, the activity is normalized according to the activity obtained for the non-exposed insects.

### Changes in midgut microbiota after exposure to indole

Indole is known to be involved in bacterial processes, either by mediating bacterial communication and quorum sensing (Kim & Park, 2013), or through antimicrobial activity (Sundar & Chang, 1993). We therefore also evaluated, in side-by-side experiments, the effect of indole exposure, as well as baculovirus infection on the larval gut microbiota load and composition. No major effect of baculovirus infection on microbiota composition and diversity was observed 48 hours posts infection. However, exposure to indole had a significant effect on the microbiota load, alpha diversity and composition (Fig 4). A multivariate canonical correspondence analysis (CCA) showed a clearly different microbial profile (P=0.012) between the indole-exposed and non-exposed group (Fig 4A). Forty-eight hours of exposure to indole, resulted in a significant decrease in gut bacterial load (P<0.019; Fig 4B) and a significant increase in bacterial diversity (P=0.03; based on the Shannon diversity index) (Fig 4C and D). The relative abundance in percentage of the top genus in each sample as depicted in Fig 4C, suggests that changes in diversity would be associated with the reduction in the relative abundance of bacteria of the genus *Enterococcus* (Fig 4C). Linear discriminant analysis effect size (LEfSE) confirms this differential abundance of the genus *Enterococcus* and revealed specific genera that were differentially enriched in each group (Fig 4E). Among the most represented genera in the indole-exposed group were *Faecalibacterium, Ruminococcus, Comanomonas*, *Chryseobacterium, Providencia*, *Sphingobium* and unclassified *Oxalobacteriaceae*, while four different genera were significantly overrepresented in the insects that were not exposed to indole. These results imply that exposure to indole changes the insect microbiota load and composition.

**Fig. 4.**
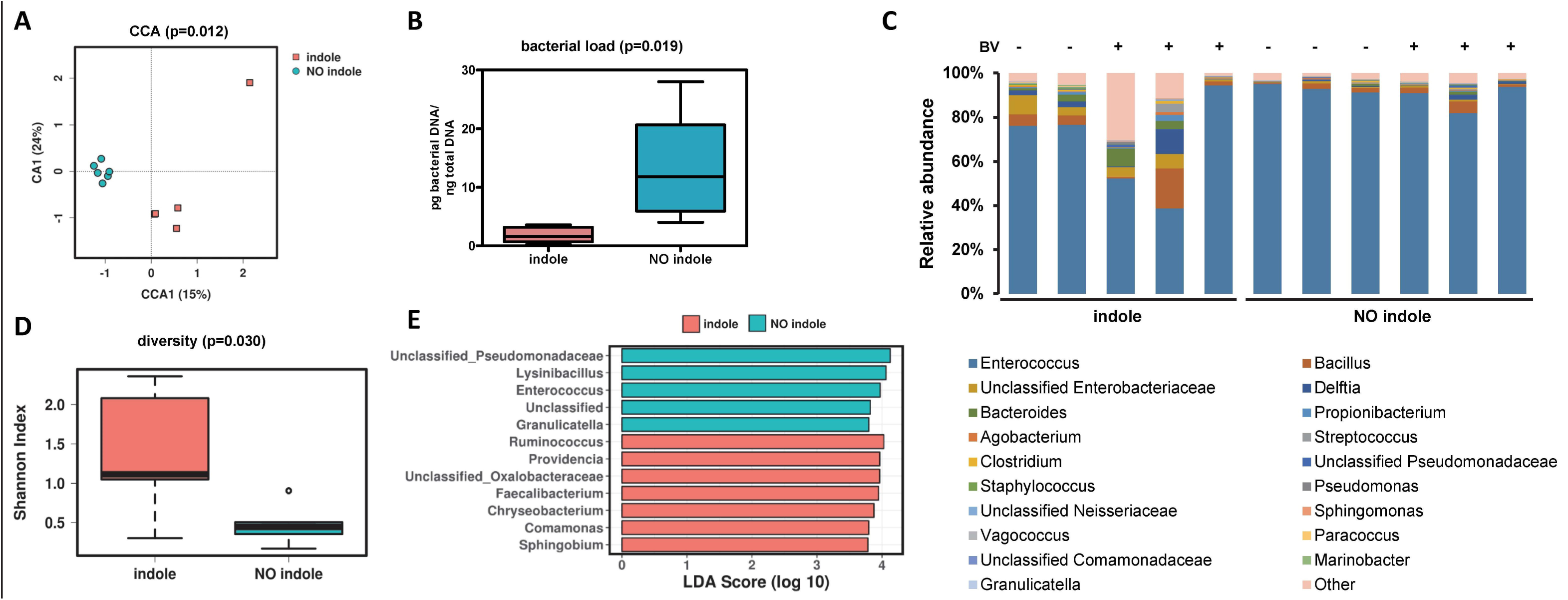
Effect of the exposure to indole on the gut microbiota composition of the *S. exigua* larvae. A) Canonical correspondence analysis (CCA) showing the relationship between gut microbiome composition (genus level) in the indole-exposed and non-exposed insects. B) Bacterial load calculated for the samples from the indole-exposed and non-exposed insects. C) Relative abundance in percentage of the top genus in samples from the indole-exposed and non-exposed insects. The exposition to the viral infection is indicated as + in the top of the panel. D) Microbial diversity calculated as the Shannon index in the samples from the indole-exposed and non-exposed insects. E) LefSe (Linear discriminant analysis effect size) results, reporting the more significantly overrepresented taxa for the indole and no indole group.

## Discussion

HIPVs play multiple roles in plant-herbivore interactions (Arimura *et al*., 2009; Maag *et al*., 2015; Turlings & Erb, 2018). Here we show that HIPVs, in addition to their already known roles, also can have a role in affecting the susceptibility of insect herbivores to viral and bacterial pathogens. These entompathogens occur naturally in the ecosystem (Hernandez-Rodriguez & Ferre, 2009; Virto *et al*., 2014; Williams *et al*., 2017), but are also used as active ingredients in biopesticides (Lacey *et al*., 2015). Specifically, we found that indole and linalool, two volatiles produced and released in response to herbivory by various plant species, have a synergistic effect on SeMNPV infectivity. To a lesser degree, the combination of indole with the bacterium *B. thuringiensis* boosted mortality caused by the bacteria in an additive manner.

In the case of indole, its effect on the susceptibility of *S. exigua* to the baculovirus was observed in different experimental settings and concentrations of the pathogen. Moreover, the synergistic mortality effect was found at a viral dose that caused only sublethal infections in most of the tested insects. At a higher viral doses, which caused mortality in most of the infected insects, the effect of indole exposure was reflected in virus virulence. This increase in virulence in the presence of indole was confirmed under more controlled conditions, where the insects were exposed under a continuous airflow and at a realistic concentration of indole.

Baculovirus infections are very common in natural populations of Lepidoptera. In the case of *S. exigua*, an important percentage of larvae in the field have a non-lethal infection of their baculovirus, SeMNPV (Virto *et al*., 2014). The dynamics of pathogen–host interactions in insects are determined primarily by host and pathogen density, but also by the virulence of the pathogen (impact on infected individuals ranging from slightly debilitating to lethal) (Myers & Cory, 2016). Our results imply that exposure to indole and linalool can increase pathogen virulence to SeMNPV to a degree that normally sublethal doses of the virus become lethal. This may have an important impact in the context of crop protection and could help to significantly decrease pest densities in the field and consequently reduce crop damage. Recent studies have started to provide evidence for selectable plant traits that enhance the ability of pathogens to control insect pests (Shikano et al., 2017). Our data further confirm the potential of plant traits to enhance the efficacy of entomopathogens as biocontrol agents. It is likely that from the extensive arsenal of metabolites produced by plants (Schwab, 2003), many others could also synergize the pest management potential of entomopathogens that are naturally found in the ecosystem or artificially released as pest control agents.

We also explored the molecular basis that underlies the effect of HIPVs on the susceptibility to entomopathogens. Indole and oxindole have previously been found to be produced by entomopathogenic bacteria and to inhibit the *in vitro* activity of PLA2, one of the key enzymes from the eicosanoids pathway that is involved in the cellular immunity (Seo *et al*., 2012; Sadekuzzaman *et al*., 2017). It has been shown that certain inhibitors of the eicosanoids pathway (including a PLA2 inhibitor), when added to the rearing diet of caterpillars (at concentrations of about 30-50 mM), can increase their susceptibility to nucleopolyhedroviruses (Stanley & Shapiro, 2009). In our study, however, when we analyzed the effect of HIPV exposure on PLA2 activity of the exposed larvae, we did not detect any reduction in the enzymatic activity for any of the three volatiles. Similarly, no effect on the PO activity, an enzyme involved in cellular and humoral defense, was observed in the insects exposed to the three HIPVs. These results imply that another mechanism, different from the direct interference with the insect’s cellular immunity, mediates the enhanced susceptibility after exposure to indole and linalool. One such mechanism could involve changes in the gut microbiota caused by the HIPVs. We and others have previously shown that changes in the gut microbiota composition can affect an insect’s susceptibility to bacterial (Broderick *et al*., 2009; Hernandez-Martinez *et al*., 2010) and viral (Jakubowska *et al*., 2013) pathogens. Insect’s gut microbiota composition and homeostasis depend on the diet (Robinson *et al*., 2010) and its immune system (Buchon *et al*., 2013), but also relies on the microbial synthesis and secretion of metabolites and enzymes that contribute to the establishment of the interactions with the host and other microbes (Lee & Hase, 2014). Gut microbiota influences in insect development and physiology (Engel & Moran, 2013), and consequently, dysbiosis in microbiota composition may have important effects on gut physiology and homeostasis leading to enhanced success of viral infections.

The changes that we observed in the gut microbial composition after indole exposure may be caused by direct effects of the indole on the microbiota, or by changes in physiological parameters of the larvae that might indirectly affect an insect’s microbiota. Given the known role of indole in microbial processes (Kim & Park, 2015), it is tempting to speculate that the observed changes are the result of direct exposure of the gut microbes to indole. More than 85 bacterial species (Gram-negative as well as gram-positive) can synthesize indole (Lee & Lee, 2010), and as an intercellular signal molecule, indole controls diverse aspects of bacterial physiology, such as spore formation, plasmid stability, drug resistance, biofilm formation, and virulence in indole-producing bacteria (Lee & Lee, 2010; Kim & Park, 2015). In our measurements species from the genus *Enterococcus* were the most dominant in the microbiota community of the *S. exigua* larvae. It is likely that indole exposure interfered with normal growth of *Enterococcus spp*, thereby probably promoting the growth of other bacterial species that could affect the insects’ physiology in a way that it lowers their resistance to entomopathogens. Vega et al (Vega *et al*., 2012) have shown that bacterial communication through indole signaling induces persistence, a phenomenon that allows a subset of an isogenic bacterial population to tolerate antibiotic treatment. Possibly, the indole-induced increase in microbial diversity occurs in a similar manner, in this case leading to enhanced susceptibility to the pathogen.

In summary, our results support a novel role for the HIPVs in the plant-insect-microbe interaction. In addition to their function in direct defence, signalling between plant tissues, and multitrophic interactions (Dicke, 2016), HIPVs may mediate interactions between insects and their pathogens. In the case of indole, probably caused by alteration of the gut microbiota composition. The observed increase in susceptibility to viral and bacterial pathogens provides an additional element to the possible application of HIPVs to regulate the abundance and dynamics of insect pests. Further experiments using other insect-pathogens combinations and other HIPVs are needed to determine the prevalence of the phenomenon and to unravel the underlying mechanisms.

## Experimental procedures

### Insects and chemicals

The *Spodoptera exigua* colony was established with eggs that were provided by Andermatt Biocontrol AG (Grossdietwil, Switzerland) and was continuously reared on artificial diet at 25 ± 3°C with 70 ± 5% relative humidity and a photoperiod of 16h light: 8h dark.

The synthetic volatiles used in the bioassays (indole, linalool and (Z)-3-hexenyl acetate) were purchased from Sigma-Aldrich.

### Effect of the HIPVs on SeMNPV infectivity

For the exposure to selected HIPVs we prepared 0.2 mL micro-centrifuge tube to which we added 4 mg of indole powder or 10 µL of 10% of linalool or 10% (Z)-3-hexenyl acetate (in destilled water). After perforating the lid of a tube with a G25 needle it was placed in a rearing well (a. 2 cm X 2 cm X 2 cm) that contained an individual larvae and a piece of artificial diet. The well was then sealed with micro perforated adhesive tape (Frontier Agricultural Sciences, Product# 9074-L).

Aiming to assess the effect of the selected HIPVs on the SeMNPV, third instar (first day) *S. exigua* larvae were orally infected and reared in presence or absence of one of the volatiles. For this, larvae were fed individually with diet plugs (about 0,4 mm^3^) containing different amounts (10^2^ or 5x10^4^) of occlusion bodies (OB) from the SeMNPV. Larvae were kept for 24 hours with the virus-contaminated food. After that, larvae that consumed completely the food were selected for the bioassay and fed with virus-free artificial diet. Larval mortality was then recorded every 12 hours until the death or pupation of all the larvae. Then mortality curves were assessed using the Kaplan-Meier method and compared using the log-rank analysis (Mentel-cox test) and the GraphPad Prism program (GraphPad software Inc., San Diego, CA, USA). In addition, and due to the different levels of mortality for each treatment, changes in virulence was estimated by comparison of the mean time to death. The statistical differences were assessed using the student’s t-test (GraphPad Prism). Three independent replicates were performed using 16 larvae per treatment and replicate.

In a second experiment, newly molted third instar larvae were exposed to the volatile indole at approximately 50 ng /h, similar to what is released by caterpillar-infested maize plants (Erb *et al*., 2015; Veyrat *et al*., 2016). For this purpose, volatile dispensers that consisted of 2 mL amber glass vials (Supelco, Sigma-Aldrich) supplied with 20 mg of synthetic indole were used. The vials were closed with an open screw cap with rubber septum. The septum were pierced with 2 µL microcaps^®^ (Drummond Scientific, Broomall, PA, USA) through which indole diffused at a constant rate. Groups of caterpillars (5 to 6) were placed in individual plastic cages (5 cm diameter, 2 cm height) covered with a nylon mesh and fed with a cube of artificial diet contaminated with 50 µL of 10^4^ OBs / mL, then kept into glass vessels which contained control or indole-releasing dispenser. Purified air entered these vessels via Teflon tubing at a rate of 0.3 L min^-1^ to avoid indole over-accumulation. The larvae were reared at 22 ± 2°C and supplied with fresh diet every 48 hours. Mortality curves and mean time to death were assessed as described above. Three independent replicates were performed using 16 larvae per treatment and replicate.

### Effect of the different HIPVs on susceptibility to B. thuringiensis infection

Effect of the selected HIPVs on the entomopathogenic bacterium *Bacillus thuringiensis* was tested using the surface contamination bioassay method (Eberle *et al*., 2012). In these experiments, a formulation of wettable granules containing *B. thuringiensis* subsp. *aizawai* (Xentari ^®^, Kenogard S.A, Spain) was tested. Surface contamination assays were employed with first instar *S. exigua* larvae, and the larvae were exposed to the different HIPVs as described in the first experiments. Briefly, a volume of 50 µL of the bacterial suspension was applied on the surface of the diet in individual wells (0.5 ng/cm^2^) and left to dry for 30-60 min in a flow hood. Then, first instar larvae were placed individually in each well together with the tube containing the respective volatile and mortality was recorded after five days. Statistical analysis were performed using either the student’s t-test or the Newman-keuls test (GraphPad Prism). Three independent replicates were performed using 16 larvae per treatment and replicate.

### Analyses of the interaction of entomopathogens with the different HIPVs

Possible antagonistic/synergistic interactions between entomopathogens and each of the selected HIPVs were determined using the mortality values at seven and five days post infection for the SeMNPV and *B. thuringiensis* treatment, respectively. Mortality percentages were corrected using the Abbott correction (Abbott, 1925). Then the expected mortality was calculated with the response addition model (Finney, 1942), which is used to evaluate mixtures of substances that have different modes of action employing the following equation:

E (c_MIX_) = E (c_A_) + E (c_B_) - [E (c_A_) * E (c_B_)], where E (c_MIX_) is the prediction of a total effect of the mixture (mortality in our case) and E (c_A_) and E (c_B_) are the observed effect caused by individual SeMNPV or *B. thuringiensis* and the volatile, respectively (Finney, 1942). Significance of the deviations between the observed and expected mortality values was assessed using Fisher’s exact test and Chi-square test.

### Effect of the HIPVs on the insect immunity

In order to study the effect of the HIPVs on the immune system of *S. exigua*, the enzymatic activities of the phenoloxidase (PO) and phospholipase A2 (PLA2), two markers of the cellular immunity, were measured. For the PO assay, hemolymph of L3 larvae exposed to a volatile or not (same conditions as above) was extracted 24 and 48h after exposure and centrifuged at 500g for 2min at 4ºC to remove the hemocytes. Four μl of cell-free hemolymph, 46μl of PBS 1X and 50μl of the substrate L-dopamine (100μg/mL in PBS 1X) were added to each wells in a 96-well microtiter plate. PO activity was determined by monitoring the increase of absorbance at 492nm for 30 min using the Infinite 200 PRO multimode plate reader (TECAN Group Ltd., Switzerland). The activity of the enzyme was represented as the initial velocity (Vo) of the reaction, measuring the change in absorbance per second. To perform the assay of PLA2 activity, bodies of the L3 larvae mentioned above were homogenized in Tris-HCl 50mM (pH 7.0) and centrifuged at maximum speed for 5 min at 4ºC. The protein concentration was determined using the Bradford (1972) assay, with bovine serum albumin (BSA) as a standard. The enzymatic reaction was done with 136μl of Tris-HCl 50mM (pH 7.0), 1μl of CaCl_2_ 150mM, 1,5μl of BSA 10%, 10μl of larval extract and 1,5μl of pyrene-labeled substrate (1-Hexadecanoyl-2-(1-Pyrenedecanoyl)-sn-Glycero-3-Phosphocholine; ThermoFisher) (10mM in ethanol). A multimode plate reader (TECAN) was used to measure fluorescence intensity by excitation at 345nm and emission at 398nm. The activity of PLA2 was then calculated as the changes in fluorescence per second. Due to the intrinsic variability between biological replicates, values for each enzyme and treatment were calculated as the difference in percentage of activity with unexposed insects within each replicate.

### Microbiota composition and diversity

To determine if exposure to indole and/or infection with the baculovirus influence the gut microbiota of *S. exigua*, third instar (first day) larvae were exposed to indole and infected with SeMNPV as described above. After 48 hours, larval midguts from each treatment were dissected, pooled by treatment and homogenized in Luria-Bertani (LB) medium supplemented with 10% of glycerol. A fraction of the homogenized guts was used for total DNA extraction using the MasterPure™ DNA Purification Kit (Epicentre, Madison, WI, USA). Three replicates were performed using 5 larvae per treatment and replicate and for each replicate the different treatments were applied simultaneously in a side-by-side manner. PCR amplification of the 16S rRNA (V3-V4 region) and sequencing were carried out using 2x 300 pb paired-end run (MiSeq Reagent kit v3) on a Illumina MiSeq sequencing platform at the Foundation for the Promotion of Health and Biomedical Research (FISABIO, Valencia). Quality assessment of obtained reads was done with the prinseq-lite program (Schmieder & Edwards, 2011) with defined parameters (i.e., min_length:50, trim_qual_right:20, trim_qual_type:mean, trim_qual_window:20). Paired reads from Illumina sequencing where joined using *fastq-join* from ea-tools suite (Aronesty, 2011). Filtered and demultiplexed sequences were then processed with the open-source software QIIME v.1.9. (Caporaso *et al*., 2010) using default parameters. A total of 12 samples was sequenced. One sequence showed fewer than 1000 reads and was removed for further analysis. The sequences were then binned into Operational Taxonomic Units (OTUs) using de novo OTU picking based on 97% identity and filtering the Unassigned taxa. Bacterial composition was also determined filtering the Unassigned, Chloroflexi and Cyanobacteria taxa, and the 20 most abundant genera were represented in a bar graphic using Excel software. Calypso version 8.2 (Zakrzewski *et al*., 2016) was used with the OTU table data transformed by CSS + log with total sum normalization, to generate Canonical Correspondence Analysis (CCA) plot for multivariate analysis at genus level, and indole exposure as factor. Alpha diversity using Shannon index and linear discriminant analysis effect size (LEfSE) (Segata *et al*., 2011) were determined at genus level, and again indole exposure as factor.

Total DNA was also used to determine the bacterial load by specific qPCR using universal primers for the 16S rDNA gene (Nadkarni *et al*., 2002). The qPCRs were carried out in StepOnePlus Real-Time PCR System (Applied Biosystems, Foster City, CA, USA). Reactions were performed using 5x HOT FIREPOL EvaGreen qPCR Mix Plus (ROX) (Solis BioDyne, Tartu, Estonia) in a total volume of 20 μl. The bacterial concentration in each sample was calculated by comparison with the Ct values obtained from a standard curve of known bacterial DNA concentration. These were generated using serial 10-fold dilutions of DNA extracted from *E. coli* bacteria. Bacterial loads were statistically compared with the student’s *t*-test (GraphPad Prism).

## Acknowledgments

This study was partially supported by the Spanish Ministry of Economy, Industry and Competitiveness and by European FEDER funds (grant AGL2014-57752-C2-2R). AF was recipient of a PhD grant from the Spanish Ministry of Education, Culture and Sport (grant FPU16/02363). We also want to thank Rosa Maria González-Martínez for her excellent help with insect rearing and laboratory management and Alejandro Tena (IVIA, Spain) for his suggestions on the data analysis.

## Competing interests

The authors declare that they have no competing interests

Fig. S1. Time course mortality of L3 larvae exposed to Indole (A), Linalool (B), and Hexenyl acetate (C) and the SeMNPV. Final mortality values are reported in Fig. 1.

Fig. S2. Time course mortality and calculated mean time to death of L3 larvae exposed to indole at a concentration of SeMNPV producing about 80% mortality. In A) and B), insects were exposed to the indole by placing 4 mg of indole in 0.2 mL tube punched with a G25 needle in their lid into the rearing well. In C) and D), insects were exposed to a continuous rate of indole (a.50 ng /h) using a volatile dispenser. * refers to P-value<0.01

